# Improved rescue of immature oocytes obtained from conventional gonadotropin stimulation cycles via human induced pluripotent stem cell-derived ovarian support cell co-culture

**DOI:** 10.1101/2023.03.27.534477

**Authors:** Alexa Giovannini, Sabrina Piechota, Maria Marchante, Kathryn S Potts, Graham Rockwell, Bruna Paulsen, Alexander D Noblett, Samantha L Estevez, Alexandra B Figueroa, Caroline Aschenberger, Dawn A Kelk, Marcy Forti, Shelby Marcinyshyn, Ferran Barrachina, Klaus Wiemer, Marta Sanchez, Pedro Belchin, Merrick Pierson Smela, Patrick R.J. Fortuna, Pranam Chatterjee, David H McCulloh, Alan Copperman, Daniel Ordonez-Perez, Joshua U Klein, Christian C Kramme

## Abstract

**Purpose:** To determine if rescue *in vitro* maturation (IVM) of human oocytes can be improved by co-culture with ovarian support cells (OSCs) derived from human induced pluripotent stem cells (hiPSCs).

**Methods:** Fertility patients undergoing conventional ovarian stimulation for oocyte cryopreservation or IVF donated denuded immature germinal vesicle (GV) and metaphase I (MI) oocytes for research, which were allocated between either the control or intervention cultures. Fertility patients aged 25 to 45 years old donated immature oocytes under informed consent, with no additional inclusion criteria. The 24-28 hour OSC-IVM culture condition was composed of 100,000 OSCs in suspension culture with human chorionic gonadotropin (hCG), recombinant follicle stimulating hormone (rFSH), androstenedione and doxycycline supplementation. The Media-IVM control lacked OSCs and contained the same supplementation. Primary endpoints consisted of MII formation rate and morphological quality assessment. Additionally, metaphase spindle assembly location and oocyte transcriptomic profiles were assessed compared to *in vivo* matured MII oocyte controls.

**Results:** We observed significant improvement in maturation outcome rates (∼1.7X) for oocytes that underwent IVM with OSCs. Specifically, the OSC-IVM group yielded a maturation rate of 62% ± 5.57% SEM versus 37% ± 8.96% SEM in the Media-IVM (*p*=0.0138, unpaired *t-*test). Oocyte morphological quality between OSC-IVM and the Media-IVM control did not significantly differ. OSC-IVM resulted in MII oocytes with no instances of spindle absence and no significant difference in position compared to *in vivo* matured IVF-MII controls. OSC-IVM treated MII oocytes display a transcriptomic signature significantly more similar to IVF-MII controls than the Media-IVM control MII oocytes did.

**Conclusion:** The novel OSC-IVM platform is an effective tool for rescue maturation of human oocytes obtained from conventional stimulation cycles, yielding oocytes with improved nuclear and cytoplasmic maturation. OSC-IVM shows broad utility for application in modern fertility treatment to improve the total number of available mature oocytes for fertility treatment.

## Introduction

Oocyte maturation is a synchronized nuclear and cytoplasmic developmental process that results in the extrusion of the first polar body (PB1) and deposition of proteins, organelles, and transcripts needed for fertilization competence and embryogenesis.^1,2^ Much of this developmental process is coordinated by several lineages of ovarian cell types including stroma, theca and granulosa cells, which send critical developmental signals to oocytes and supporting cells through paracrine and autocrine mechanisms.^3,4^

The current standard of care for *in vitro* fertilization (IVF) treatment is to discard all immature oocytes shortly following retrieval, imposing a key early obstacle to maximizing the final yield of useable oocytes.^5^ For some patients, many of the oocytes obtained using standard stimulation are immature at retrieval, limiting the number of usable oocytes for freezing or transfer.^6^ Further, for patients with diminished ovarian reserve, the total oocyte number retrieved is small which makes every oocyte highly valuable for cycle success. Therefore, in IVF and in particular for patients of advanced age, patients with a low total oocyte retrieval number, or patients with a large number of immature oocytes, effective rescue *in vitro* maturation (IVM) holds great value as a strategy to maximize the use of all oocytes retrieved during the IVF process.

Although immature oocytes removed from the follicular environment can undergo meiosis, their rate of maturation and subsequent developmental competence has not been sufficiently reliable for widespread clinical use.^1,7^ Decades of research into how oocytes develop have led to the creation of IVM-stimulating cell culture media, which are commercially available for human oocyte IVM. Such IVM media products are usually designed for oocytes enclosed in cumulus cells, limiting their utility for denuded immature oocyte rescue applications.^8–19^ However, in most traditional fertility treatment settings, oocytes are denuded of cumulus cells upon retrieval in preparation for intracytoplasmic sperm injection (ICSI) or oocyte cryopreservation, and the denuded immature oocytes either containing a germinal vesicle (GV) or lacking a germinal vesicle and polar body (metaphase I, MI) are left with no alternative use other than being discarded.^20^ Numerous studies have supported that oocytes denuded of cumulus cells are capable of maturation, and their rate of maturation can be improved through supplementation of various growth factors, cumulus cells, or granulosa cells.^5,21–25^ While the precise mechanism of improving denuded oocyte maturation rates is often unknown, it is likely that paracrine signaling and regulation of the local nutrient environment can influence oocyte meiotic resumption and completion.^5,26^ Indeed, as a consequence of standard clinical practice in ICSI of removing the oocyte’s natural cumulus support cells, methods that replace functions of those support cells stand to possibly improve oocyte maturation rate and quality. Therefore, new approaches to IVM that include alternatives for denuded immature oocytes are necessary to maximize oocyte usage and ensure comparable developmental quality to mature oocytes used in IVF in a robust and reproducible manner. This advance will allow expanded application of IVM for denuded oocytes retrieved in standard of care cycles.

Our group has recently demonstrated a novel technology to generate ovarian support cells (OSCs) from human induced pluripotent stem cells (hiPSCs) in a rapid, efficient, and reproducible manner through transcription factor (TF)-directed differentiation.^27^ The OSCs are primarily composed of FOXL2+ AMHR2+ NR2F2+ granulosa-like cells. These cells produce growth factors and are steroidogenic in the presence of follicle stimulating hormone (FSH). In this study, we investigate the potential of hiPSC-derived OSCs to rescue immature denuded oocytes retrieved in conventional controlled ovarian hyperstimulation cycles and assess their morphological and transcriptomic quality compared to *in vivo* and *in vitro* matured control oocytes.

## Materials and Methods

### Collection of Immature Oocytes

#### Subject ages, IRB, and informed consent

47 fertility patient subjects undergoing treatment at Extend Fertility New York and Ruber Clinic Madrid were enrolled in the study for donating immature oocytes using informed consent (IRB# 20222213, Western IRB). Subject ages were between 25 and 45 years of age, with an average age of 35. An additional cohort of 34 metaphase II (MII) oocytes obtained from conventional controlled ovarian stimulation (COS) were donated from RMA Clinic of New York for transcriptomic analysis and served as reference controls (IRB#20222213).

#### Oocyte retrieval and donation

Fertility patients providing discarded immature oocytes had signed informed consents, provided by the clinic, permitting their use for research purposes. Patients underwent typical age-appropriate controlled ovarian stimulation using gonadotropin releasing hormone (GnRH) analogs (agonist or antagonist) and injections with recombinant or highly purified urinary gonadotropins (recombinant FSH, human menopausal gonadotropins) followed by an ovulatory trigger (human Chorionic Gonadotropin (hCG) or GnRH agonist). 34-36 hours following the trigger injection(s), oocytes were retrieved from the patient under conscious sedation using standard clinical procedures.

Retrieved oocytes were exposed to hyaluronidase briefly then adherent cumulus cells were mechanically removed by repeatedly drawing up and expelling each cumulus-oocyte complex with a small-bore pipette. Denuded oocytes were assessed for maturation by observation of a polar body and/or a germinal vesicle. Immature oocytes (GV or MI), which would usually be discarded, were instead allocated to the research study. All immature oocytes retrieved from the clinic each day were pooled and placed in gassed LAG Medium (Medicult, Cooper Surgical) in a 5ml round-bottom tube that was transferred from the clinic to the research laboratory in a 37°C transport incubator.

For some experiments, immature (GV and MI) oocytes from similar ICSI and egg freezing cycles were vitrified and stored at the clinics. Cryopreserved oocytes were transported from the clinic to our laboratory in liquid nitrogen and stored until use. Oocytes were then thawed using the standard Kitazato protocol for vitrified or standard Vitrolife protocol for slow frozen oocytes (Kitazato, USA and Vitrolife, USA), evaluated for maturation status as GV or MI, and used for comparisons of *in vitro* maturation conditions.

A limited number of MII oocytes obtained from conventional controlled ovarian stimulation, which were previously banked for research purposes, were provided as controls for this study (IVF-MII). These oocytes were transferred to our laboratory and thawed using either the standard Kitazato protocol for vitrified oocytes (Kitazato, USA) or slow freeze-thaw protocol for previously slow frozen oocytes (Vitrolife, USA), and utilized for live fluorescent imaging and transcriptomic analysis.

### Preparation of Ovarian Supporting Cells (OSCs)

Human induced pluripotent stem cell (hiPSC) derived OSCs were created according to transcription factor (TF)-directed protocols described previously.^27^ Briefly, OSCs were produced through monolayer directed-differentiation using the transcription factors *NR5A1, RUNX2*, and *GATA4* in a five day differentiation protocol from a single monoclonal hiPSC line described in a previous study.^27^ OSCs were produced in multiple batches and cryopreserved in vials of 120,000 to 150,000 cells each, and stored in the vapor phase of liquid nitrogen in CryoStor CS10 Cell Freezing Medium (StemCell Technologies). OSC batches were confirmed FSH-tuned steroidogenic through estradiol and progesterone ELISA assays and for presence of key granulosa cell markers FOXL2, CD82, and lack of hiPSC markers OCT4 and TRA-160 via flow cytometry.

Culture dishes (4+8 Well Dishes, BIRR) for oocyte maturation experiments were prepared with culture media and additional constituents in 100μl droplets under mineral oil (LifeGuard, LifeGlobal Group) the day before oocyte collection and equilibrated in the incubator overnight. The morning of oocyte collection, cryopreserved OSCs were thawed for 2-3 minutes at 37°C (in a heated bead or water bath), resuspended in OSC-IVM medium and washed twice using centrifugation cell pelleting to remove residual cryoprotectant. Equilibrated OSC-IVM medium was used for final cell resuspension. OSCs were then plated at a concentration of 100,000 OSCs per 100μl droplet by replacing 50μl of the droplet with 50μl of the OSC suspension 2-4 hours before the addition of oocytes to allow for culture equilibration and culture media conditioning (Figure 1A). OSCs were cultured in suspension culture surrounding the denuded oocytes in the microdroplet under oil. IVM culture proceeded for 24 to 28 hours, after which oocytes were removed from culture, imaged, and flash frozen for molecular analysis.

**Figure 1:**
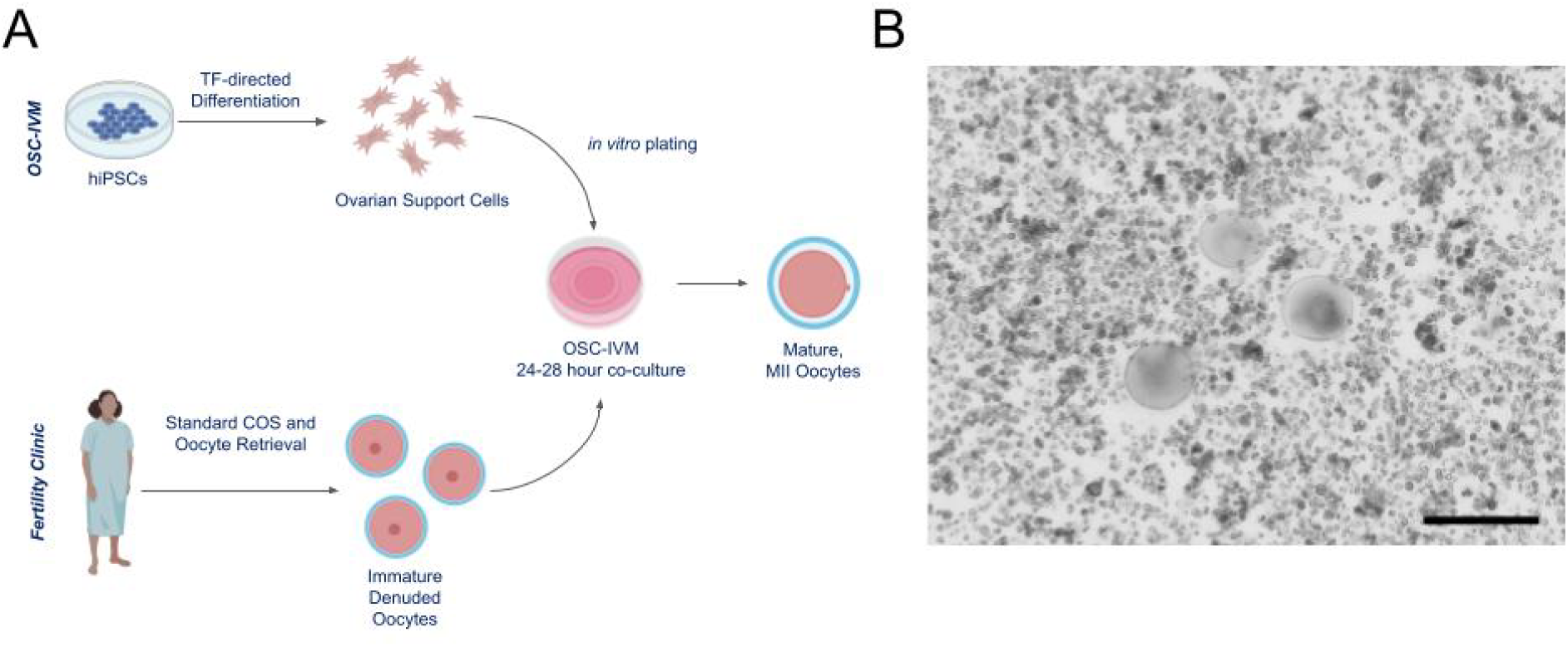
Ovarian support cells (OSC) rescue IVM co-culture system. **A** Schematic of the experimental co-culture IVM approach. hiPSCs were differentiated using inducible transcription factor overexpression to form ovarian support cells (OSCs). Human oocytes were obtained from patients in the clinic after standard gonadotropin stimulation, and immature oocytes (GV and MI) identified after denuding were allocated to this research study. In the embryology lab, dishes were prepared including OSCs seeding as required, and immature oocytes were introduced for IVM co-culture. Oocyte maturation and health were assessed after 24-28 hours IVM co-culture, and oocytes were banked for further analyses. **B** Representative image of co-culture containing immature human oocytes (n=3) and OSCs. Scale bar: 200μm. Denuded GV oocytes are seen with surrounding OSCs in suspension culture.

### In Vitro Maturation

Immature oocytes were maintained in preincubation LAG Medium (Medicult, Cooper Surgical) at 37° C for 2-3 hours after cumulus removal prior to introduction to *in vitro* maturation conditions (either Media-IVM or OSC-IVM). Commercially-available Medicult IVM Medium (Medicult, Cooper Surgical) was used as the base media for our studies. Although this medium was designed for *in vitro* maturation of cumulus-enclosed oocytes, there are no commercially-available media expressly designed for use with denuded oocytes following a retrieval employing an LH surge-mimicking trigger. Therefore, we have used media that we believe are the best available option for examining rescue *in vitro* maturation of denuded immature oocytes.

#### Experiment (OSC activity)

The purpose of this comparison was to determine whether the stimulated OSCs were the active ingredient or driver of oocyte maturation in the IVM co-culture system. In both the Experimental and Control conditions, media were prepared by following the manufacturer’s recommendations, and further supplemented with androstenedione and doxycycline (both necessary for activation of OSCs) in order to compare maturation outcomes with or without OSCs in the same medium formulation (see Table 1).

#### Oocyte culture condition description

Donated oocytes were retrieved from 47 patients and pooled into 29 independent cultures, totaling 141 oocytes with 82 utilized in OSC-IVM (Experimental) and 59 utilized in Media-IVM (Control). Due to low and highly variable numbers of discarded immature oocytes acquired per donation, oocytes from each donor pool were distributed equitably between the two conditions when possible or into one condition at a time, with no more than 15 oocytes per culture well. *In vitro* maturation occurred at 37°C for 24-28 hours in a tri-gas incubator with CO_2_ adjusted so that the pH of the bicarbonate-buffered medium was 7.2-7.4 and with the O_2_ level was maintained at 5%.

### Assessment of oocyte *in vitro* maturation

At the end of the *in vitro* culture, oocytes were harvested, mechanically denuded and washed of any residual OSCs. Oocytes were then individually assessed for maturation state according to the following criteria:

**GV** - presence of a germinal vesicle, typically containing a single nucleolus within the oocyte.

**MI** - absence of a germinal vesicle within the oocyte and absence of a polar body in the perivitelline space between the oocyte and the zona pellucida.

**MII** - absence of a germinal vesicle within the oocyte and presence of a polar body in the perivitelline space between the oocyte and the zona pellucida.

Following assessment of *in vitro* maturation and morphology scoring, oocytes were individually imaged using digital photomicrography and if required, examined by fluorescent imaging for the second meiotic metaphase spindle. No oocytes from this study were utilized or banked for fertilization, transfer, implantation, or reproductive purposes.

### Oocyte morphology scoring

Oocytes harvested post-IVM were individually imaged using digital photomicrography on the ECHO Revolve inverted fluorescent microscope using phase contrast imaging. The images were later scored according to the Total Oocyte Score (TOS) grading system.^28^ A single trained embryologist was blinded and oocytes were given a score of -1, 0, 1 for each of the following criteria: morphology, cytoplasmic granularity, perivitelline space (PVS), zona pellucida (ZP) size, polar body (PB) size, and oocyte diameter. ZP and oocyte diameter were measured using ECHO Revolve Microscope software and the image analysis software FIJI (2.9.0/1.53t). The sum of all categories was taken to give each oocyte a total quality score, ranging from -6 to +6, with higher scores indicating better morphological quality.

### Examination of the second meiotic metaphase spindle and its position relative to the polar body

Previously vitrified denuded immature oocytes were thawed using the standard Kitazato protocol (Kitazato, USA) and equitably distributed across OSC-IVM and Media-IVM conditions for 28 hour cultures. Additional donated MII oocytes were collected and live cell stained to visualize the microtubules of the meiotic spindle apparatus by fluorescent microscopy as an IVF control (IVF-MII) (Figure 3). MII oocytes were incubated in 2μM of alpha-tubulin dye (Abberior Live AF610) for one hour in the presence of 10μM verapamil (Abberior Live AF610). Spindle position was then visualized using fluorescence microscopy (ECHO Revolve microscope, TxRED filter block EX:560/40 EM:630/75 DM:585). The angle of the first polar body and spindle apparatus in the IVM oocytes was determined (with the vertex at the center of the oocyte) using FIJI software.^29^ This measurement was also made on the cohort of IVF-MII oocytes as a control reference population. Oocytes were handled minimally and immediately imaged under epi-fluorescent microscopy to minimize disturbance to oocyte health and quality during live imaging.

### Cryopreservation of oocytes for subsequent molecular analyses

Following the completion of morphological examination, oocytes were individually placed in 0.2ml tubes containing 5μl Dulbecco’s Phosphate Buffered Saline (DPBS) and flash frozen in liquid nitrogen, and were stored at -80°C until subsequent molecular analysis.

### Single oocyte transcriptomics library preparation and RNA sequencing

Libraries for RNA sequencing were generated using the NEBNext Single Cell/Low Input RNA Library Prep Kit for Illumina (NEB #E6420) in conjunction with NEBNext Multiplex Oligos for Illumina (96 Unique Dual Index Primer Pairs) (NEB #E6440S), according to the manufacturer’s instructions. Briefly, oocytes frozen in 5μl DPBS and stored at -80°C were thawed and lysed in NEB Lysis Buffer, then RNA was processed for reverse transcriptase and template switching. cDNA was PCR amplified with 12-18 cycles, then size purified with KAPA Pure Beads (Roche). cDNA input was normalized across samples. Following fragmentation and end prep, NEBNext Unique Dual Index Primer Pair adapters were ligated, and samples were enriched using 8 cycles of PCR. Libraries were cleaned up with KAPA Pure Beads, quantified using Quant-iT PicoGreen dsDNA Assay Kit (Invitrogen), then an equal amount of cDNA was pooled from each oocyte library. The pool was subjected to a final KAPA Pure bead size selection if required and quantified using Qubit dsDNA HS kit (Invitrogen). After verification of library size distribution (∼325bp peak) using Bioanalyzer HS DNA Kit (Agilent), the library pool was subjected to RNA sequencing analysis using the MiSeq Micro V2 (2x150bp) or MiSeq V2 (2x150bp) kit on an Illumina MiSeq according to the manufacturer’s instructions.

### Oocyte transcriptomics data analysis

Illumina sequencing files (bcl-files) were converted into fastq read files using Illumina bcl2fastq (v2.20) software deployed through BaseSpace using standard parameters for low input RNA-seq of individual oocytes. Low input RNA-seq data gene transcript counts were aligned to *Homo sapiens* GRCH38 (v2.7.4a) genome using STAR (v2.7.10a) to generate gene count files and annotated using ENSEMBL.^30^ Gene counts were combined into sample gene matrix files (h5). Computational analysis was performed using data structures and methods from the Scanpy (v1.9.1) package as a basis. Gene transcript counts were normalized to 10,000 per sample and log (ln) plus 1 transformed. Principal component analysis was performed using Scanpy package methods focusing on 30 PCA components. Integration and project (batch) correction was performed using BBKNN.^31^ Projection into two dimensions was performed using the Uniform Manifold Approximation and Projection (UMAP) method.^32^ Cluster discovery was performed with the Ledien methods with resolution of 0.5.^33^

To define the expected transcriptomic profile for normal MII oocytes we used the donated cohort of *in vivo* matured IVF-MII samples (n=34) as a reference point and compared this to subsets of the post-IVM GV cells using differential gene expression. The top 50 differentially expressed genes were collected for each comparison using both the Wilcoxon ranked sum test and the cosine similarity-based marker gene identification (COSG) method.^34^ No other MI or MII oocyte sets were used as reference points, as these marker genes were developed to ensure minimal bias for other MII transcriptomic profiling. This method generated the ‘failed-to-mature GV’ and ‘IVF-MII’ signature marker gene expression profiles. Cells were scored for similarity to each marker gene set using Scanpy gene marker scoring methods.

To visualize our cells in signature marker score space, we plotted the marker scores in two dimensions. We then manually divided the space into quadrants based on morphological maturation outcomes and Leiden clusters. Clusters were annotated taking into consideration their distribution in signature marker score space and presence in each quadrant correlating their IVM maturation outcome and whole transcriptomic profiles.

### Data analysis and statistics

Oocyte maturation outcome data was analyzed using Python statistical packages pandas (1.5.0), scipy (1.7.3), and statsmodels (0.13.2). Maturation percentages by donor group were analyzed using unpaired *t-*test as functions of the IVM environment as OSC-IVM or Media-IVM. *t*-test statistics were computed comparing OSC-IVM versus Media-IVM, then used to calculate *p*-values using Welch’s correction for unequal variance. One-way ANOVA was utilized for comparisons of more than two groups for spindle apparatus location analysis. Chi-squared analysis was utilized for comparison of the Leiden group population make up in transcriptomic analysis for the three sample conditions. Bar graphs depict mean values for each population and error bars represent standard error of the mean (SEM). Number of independent oocytes for the experiment are indicated in the figure and Materials and Methods.

## Results

### hiPSC-derived OSCs effectively promote human oocyte maturation in co-culture with denuded oocytes

We have previously demonstrated that hiPSC-derived OSCs are predominantly composed of granulosa-like cells.^27^ In response to FSH stimulation *in vitro*, the OSCs produce growth factors and steroids necessary for paracrine interaction with oocytes and cumulus cells.^27^ To investigate whether hiPSC-derived OSCs are functionally capable of promoting human oocyte maturation *in vitro*, as an approach to rescue immature denuded oocytes, we established a co-culture system of these cells with retrieved denuded immature oocytes and assessed maturation rates after 24-28 hours (Figure 1).

We first examined whether OSC-IVM affected the rate of maturation of denuded oocytes compared to oocytes in the Media-IVM control containing the same culture medium and all supplements but no OSCs. Strikingly, we observed significant improvement in maturation outcome rates (∼1.7X) for oocytes that underwent rescue-IVM with OSCs. Specifically, OSC-IVM yielded an oocyte maturation rate of 62% ± 5.57% SEM versus 37% ± 8.96% SEM in the Media-IVM (Figure 2A, *p*=0.0138, unpaired *t-*test). We additionally scored the morphological quality of MII oocytes obtained in both rescue-IVM conditions by assessing the Total Oocyte Score (TOS). We found no significant difference between the two groups (Figure 2B, *p*=0.5725, unpaired *t-*test), suggesting that *in vitro* maturation of denuded oocytes does not affect the morphological features of MIIs. Altogether, these data indicate that OSC co-culture increases the rate of oocyte maturation compared to spontaneous maturation observed in the control IVM media, without any detrimental effect on human oocyte morphological quality. This finding highlights the potential for the use of hiPSC-derived OSCs for rescuing immature denuded oocytes from IVF procedures.

**Figure 2:**
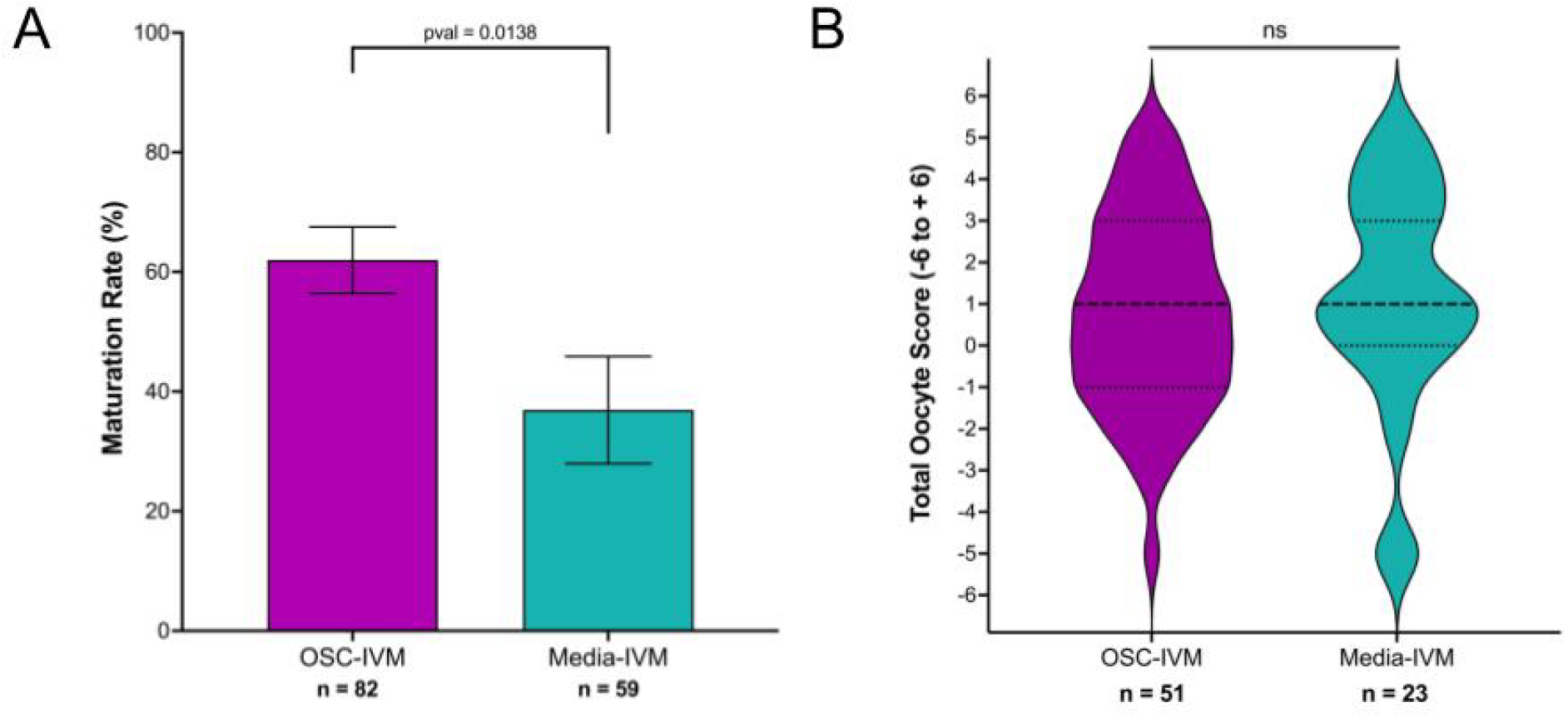
OSCs improve human oocyte rescue maturation rates compared to IVM medium lacking OSCs. **A** Maturation rate of oocytes after 24-28 hour IVM experiments, including oocyte co-culture with OSCs (OSC-IVM), or in Media Control (Media-IVM). *n* indicates the number of individual oocytes in each culture condition. Error bars indicate mean ± SEM. *p*-value derived from unpaired *t*-test comparing Experimental OSC-IVM to Control Media-IVM. Due to low numbers of retrieved oocytes per donor, each group contains oocytes from predominantly non-overlapping donor groups and pairwise comparisons are not utilized. **B** Total Oocyte Scores (TOS) generated from imaging analysis of MII oocytes after 24-28 hour IVM experiments. *n* indicates the number of individual MII oocytes analyzed. Median (dashed lines) and quartiles (dotted lines) are indicated. Unpaired *t-*test indicated no significant (ns, *p*=0.5725) difference between the means.

### OSC-IVM promotes high quality assembly of the second meiotic spindle apparatus in rescue-IVM oocytes

Second meiotic spindle assembly, more specifically both the presence of and the angle of the spindle relative to PB1, has been implicated in previous studies as a key indicator of oocyte quality relevant to fertilization and developmental competence, with a smaller angle indicating improved quality.^29^ We sought to determine the relative position of the second meiotic spindle apparatus and PB1 in OSC-treated oocytes in comparison to MII oocytes retrieved from IVF cycles (IVF-MII), as a measure of oocyte quality (Figure 3). Due to the desire to perform live cell imaging, with minimal deleterious handling of the oocytes, fixation and confocal microscopy of the oocytes was avoided in favor of live cell imaging dyes. We also included as a control the oocytes that matured in the Media-IVM condition (Figure 3). We found that the spindle angle was not significantly different between conditions (MII OSC-IVM: 22° ± 5.2 SEM; MII Media-IVM: 15° ± 5.7 SEM; IVF-MII: 41° ± 8.3 SEM; *p*=0.1155; ANOVA), suggesting that *in vitro* maturation of denuded oocytes does not impair spindle position. Interestingly, the only condition in which we did not observe instances of spindle absence was MII oocytes derived from OSC-IVM (Figure 3B). More studies are needed to validate the relevance of this observation, but it is likely to indicate the formation of high-quality oocytes. Altogether, these results indicate that MII oocytes matured *in vitro* in combination with OSCs hold equivalent spindle angle values to MII oocytes directly retrieved from IVF procedures, suggesting that rescue-IVM applied to rescue denuded immature oocytes is not detrimental to oocyte quality based on this parameter.

**Figure 3:**
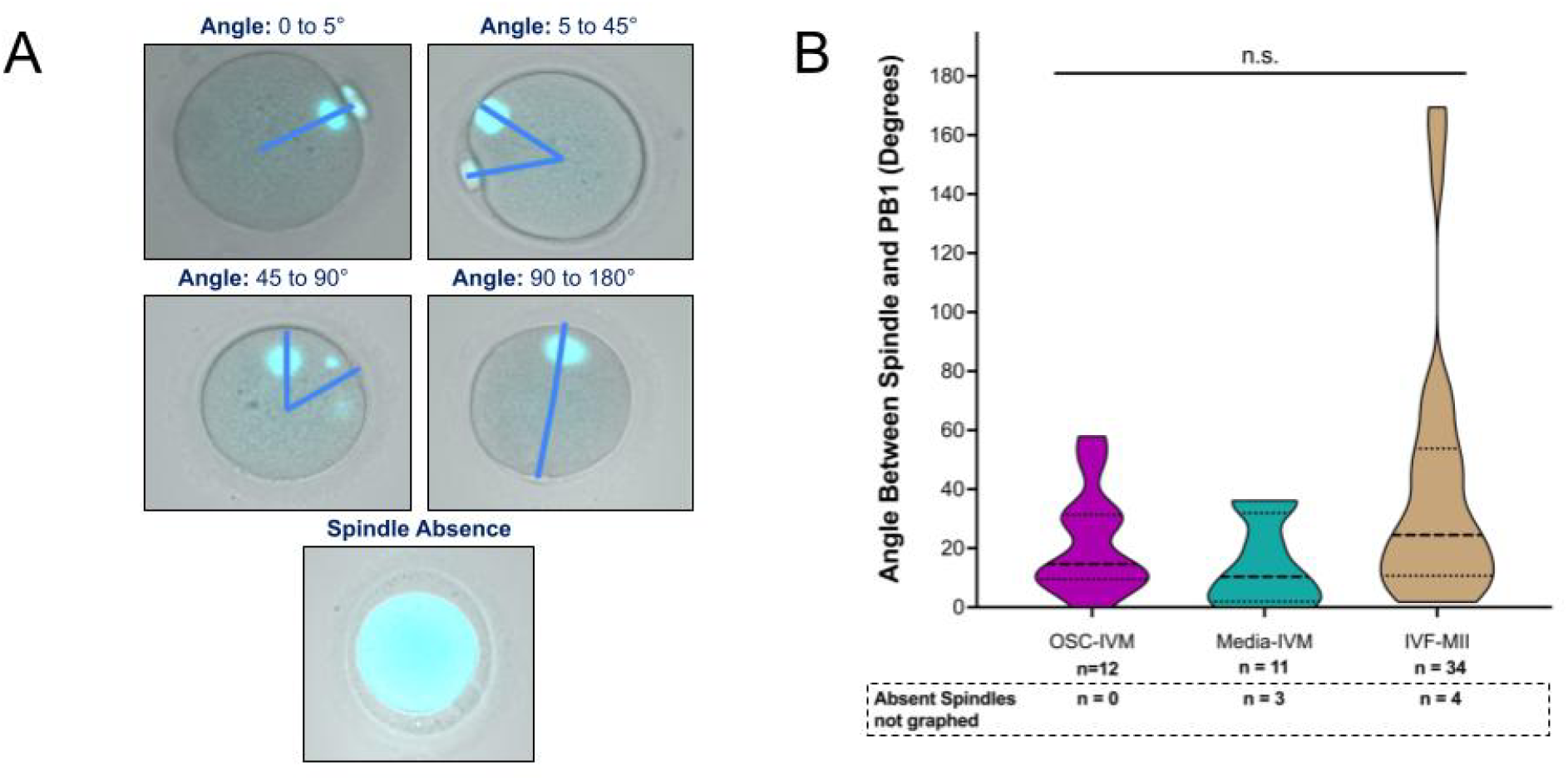
OSC-IVM oocytes effectively assemble the second meiotic spindle apparatus. **A** Representative images of MII oocytes after 28 hour IVM co-culture with OSCs, stained with fluorescent alpha-tubulin dye (cyan) to visualize the meiotic spindle. Blue lines transecting the middle of the PB1 and the spindle apparatus from the oocyte center were used to derive the PB1-spindle angle. PB1-spindle angle ranges are indicated above. An example of an MII with a missing spindle apparatus is provided from the Media-IVM condition. **B** Quantification of the angle between the PB1 and spindle apparatus, derived from oocyte fluorescence imaging analysis (as in **A**). *n* indicates the number of individual oocytes analyzed from each condition. Number of MII oocytes with no spindle assembly observed is also indicated below the axis labels in the dashed box. Median (dashed line) and quartiles (dotted line) are indicated. ANOVA statistical analysis found no significant difference (ns, *p*=0.1155) between the means of each condition.

### OSC-IVM promotes maturation of MII oocytes with high transcriptomic similarity compared to *in vivo* matured MII oocytes

To further compare the quality and maturation of OSC-IVM oocytes relative to a cohort of IVF-MII control oocytes and the Media-IVM oocytes, we performed single oocyte transcriptomic analysis. Transcriptomic analyses provide a global view of oocyte gene expression, providing a strong representation of their cellular state, function, and general attributes.^35–37^ We started by combining our transcriptomic datasets that included: 1) denuded immature oocytes after 24-28 hours in co-culture with OSCs (OSC-IVM); 2) denuded immature oocytes cultured in the *in vitro* maturation media control (Media-IVM); and, 3) MII oocytes retrieved from regular COS-IVF cycles (IVF-MII). We generated UMAP plots and annotated individual oocytes by Condition (OSC-IVM, Media-IVM, and IVF-MII) and Maturation outcome (GV, MI, MII) (Figure 4A). From this analysis, we observed that maturation state was the main driver of oocyte separation in whole transcriptomic space, suggesting that transcriptional profiles are a good predictor of oocyte maturation state. MII oocytes project predominantly into the large cluster on the upper right of the UMAP plot (Figure 4A Maturation), while GV oocytes project predominantly into a smaller cluster on the lower left of the UMAP plot. Hence, the separation in the UMAP is a combination of the two projected dimensions. In this UMAP representation, MII oocytes retrieved from IVF (IVF-MII) show close grouping together with MII from both the OSC-IVM, as well as Media-IVM. Similarly, GVs from OSC-IVM and Media-IVM were closely located and distinct from the MII oocytes. In contrast, MI oocytes were scattered across both groups, a likely consequence of their intermediate maturation state and being present in very low numbers in comparison with the other two populations (GVs and MIIs).

**Figure 4:**
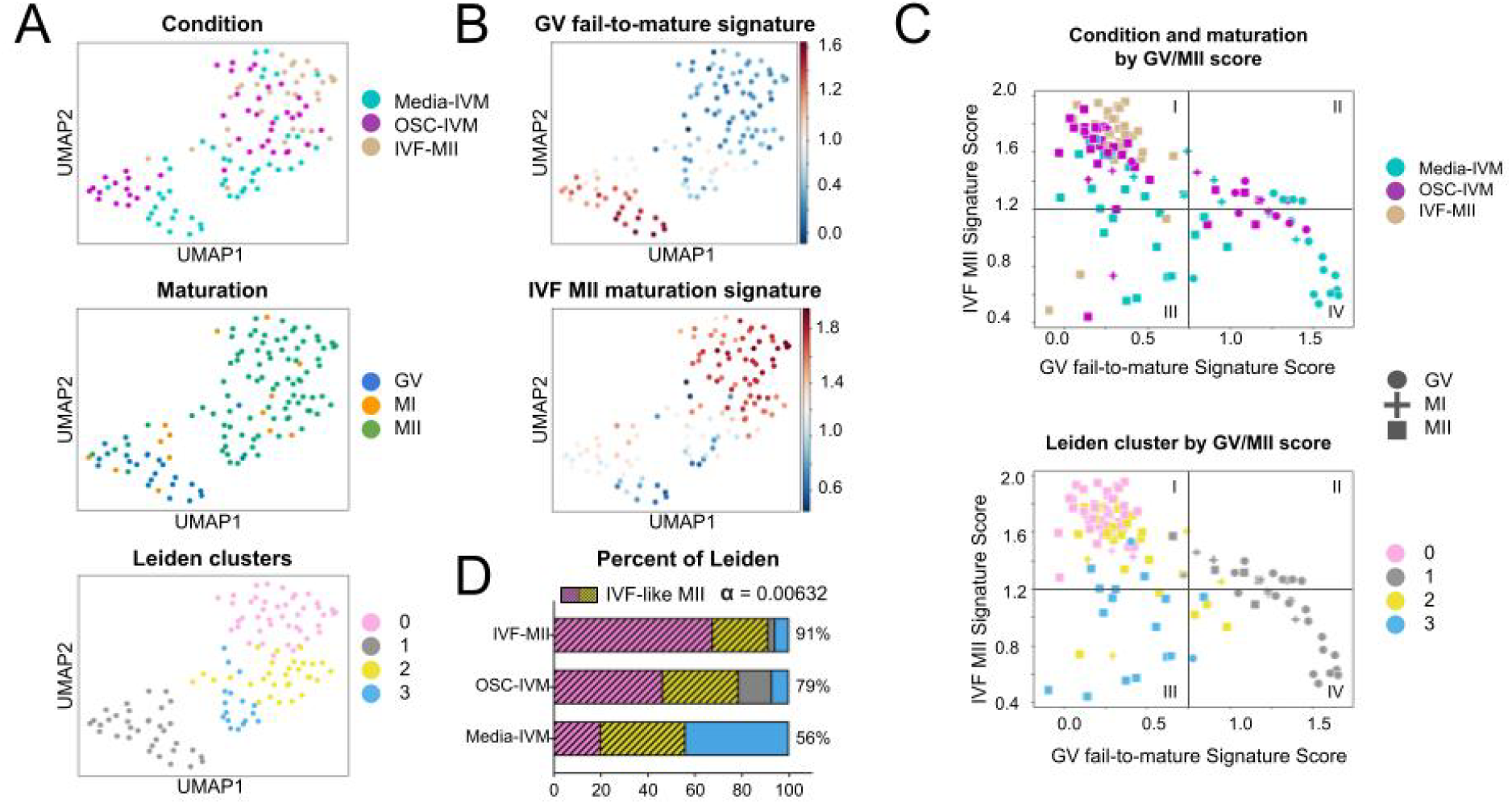
OSC-IVM oocytes share transcriptomic similarity with IVF MII oocytes. **A** UMAP projections of oocyte transcriptomes with symbols colored by experimental condition (OSC-IVM, Media-IVM, IVF-MII), oocyte maturation state, and Leiden cluster. Each symbol represents one oocyte. *n*=81 oocytes. **B** UMAP projections colored by scores for each of the gene marker sets (GV fail-to-mature and IVF-MII). **C** UMAP projection generated from the scores of cells for each of the two signature marker sets (GV vs IVF-MII), colored by experimental condition, oocyte maturation state, and Leiden cluster. **D** Quantification of MII oocytes belonging to the “IVF-like” clusters (0,2) by experimental condition (OSC-IVM, Media-IVM, or IVF-MII), with color distribution indicating percentage of population in each Leiden cluster. Striped bars are utilized to denote clusters with predominantly IVF-like characteristics.

To transcriptomically assess our rescue-IVM oocytes, we generated reference transcriptomic signatures for oocyte maturation outcomes. We used MII oocytes retrieved from conventional ovarian stimulation IVF sample (IVF-MII) to create a gene score for *IVF MII maturation signature*. In parallel, we used the stalled GVs resultant from rescue-IVM conditions (OSC-IVM and Media-IVM) to generate a gene score for *GV fail-to-mature signature* (Figure 4B). These two gene signatures were utilized to capture a relative positive control, namely an IVF-like successful maturation outcome, as well as a negative control, namely oocytes that arrest as GVs.

To better understand transcriptomic nuances amongst the mature MII oocytes, we used the Leiden algorithm to further subcluster our samples into groups sharing closer transcriptomic profiles. We identified three clusters (0, 2, and 3) within the MII oocytes population, and one cluster (1) comprised mostly GVs. As expected, the GV maturation signature was strongly represented in cluster 1. Similarly, the MII maturation signature included MIIs from both IVF and rescue-IVM, and IVF MIIs were more overrepresented in clusters 0 and 2 compared to cluster 3. As such, we designated cluster 1 as representing the (GV) failed maturation transcriptomic profile, while clusters 0 and 2 represented a profile similar to the IVF MII maturation transcriptomic profile. Interestingly, cluster 3 showed lower expression for both the IVF MII and rescue-IVM GV failed maturation signatures. This could indicate a transitional state between immature and mature oocytes in which neither signature is highly upregulated, or could result from cell activity stasis, shutdown, or oocyte stalling.

In Figure 4C, we assess the quality of individual oocytes relative to the *IVF MII maturation signature* (y-axis), as well as the *GV fail-to-mature signature* (x-axis). For visual clarity we divide our signature dimension plot into quadrants to help denote the separation between classification groups. As expected, we observe that most of the oocytes morphologically classified as GVs clustered in the lower right quadrant (IV), holding a high score for *GV fail-to-mature signature* along with a low score for *IVF MII maturation signature*. In contrast, individual oocytes from the IVF-MII condition clustered together (∼91%) in the upper left quadrant (I), with a high score for *IVF MII maturation signature* and a low score for *GV fail-to-mature signature*. Strikingly, OSC-IVM MIIs were found mostly (∼79%) in the upper left quadrant (I) along with the IVF-MII oocytes, suggesting a strong transcriptomic similarity between these two groups. In contrast, MIIs from the Media-IVM were often (∼44%) located on the lower left quadrant (III) depicting a low score for both *IVF MII maturation signature* and *GV fail-to-mature signature*. Interestingly, the lower left quadrant (III) predominantly comprised cells derived from cluster 3, which despite their weak *MII maturation signature*, were morphologically classified as MIIs. This divergence in morphological classification and transcriptomic profile suggests that these oocytes are possibly in an incomplete transitional phase of maturation and that the cytoplasmic maturity is not synchronized with the nuclear maturity. To assess for confounding variables in our transcriptomic analysis, we evaluated expression of cell cycle, apoptosis and oxidative stress genes and did not detect any significant patterns, indicating that the oocytes do not exhibit major signs of stress or breakdown (Supplemental Figure 1). Altogether this dataset suggests that MII oocytes derived from OSC-IVM were transcriptionally more similar to those from the IVF-MII condition than Media-IVM control MII oocytes were.

Finally, to determine the ratio of MII oocytes with a strong *IVF MII maturation signature* in each experimental condition, we calculated the percentage of cells in clusters 0 and 2, identified as containing oocytes with a ‘IVF-like MII’ signature (Figure 4D). As expected, the strong majority (91%) of MII oocytes from the IVF-MII condition were classified within clusters 0 and 2. As a continued indication of positive maturation impact, co-culture with OSCs led to generation of 79% MII oocytes with a ‘IVF-like MII’ profile (cluster 0 and 2). In comparison, just 56% of resultant MII oocytes from the Media-IVM condition were found in ‘IVF-like MII’ profile clusters. This population distribution is significantly different from random (χ^2^ test, *α* = 0.00632). Altogether, we conclude that OSC-IVM supports formation of MII oocytes with high transcriptomic similarity to IVF matured MII oocytes and highlights the potential of using this novel approach to rescue denuded immature oocytes from IVF procedures.

## Discussion

In this study, we demonstrate the utilization of human stem cell-derived OSCs for rescue *in vitro* maturation of human denuded oocytes. These results demonstrate that OSC co-culture may be an effective platform for rescue maturation of immature denuded oocytes following conventional follicular stimulation by recreating their support cell environment that is lost during denuding. Our work is the first of its kind to explore the potential of allogeneic stem cell-derived OSCs as a tool for rescue IVM of denuded oocytes in humans and shows the value of this and other cell engineering approaches for improving IVF treatments and patient success outcomes.

It is generally a concern that IVM and rescue-IVM can result in oocytes of poor morphological quality and with gene expression profiles that are significantly different from conventional COS oocytes.^38^ Our measures of morphology indicate that there is no striking difference in the morphological quality between rescue-IVM conditions and no significant difference in the angle between the spindle and PB1 compared to IVF MIIs. However, it should be noted that the angle between the spindle and PB1 in the IVF group could be broadened because the already-present PB1 can be displaced during removal of the cumulus, whereas this does not occur when cumulus is removed from oocytes prior culture for rescue-IVM and prior to PB1 extrusion. Our analysis additionally showed OSC-IVM MII oocytes had gene expression signatures that were more similar to IVF-MII controls than MII oocytes from Media-IVM oocytes had compared to IVF-MII controls. Based on this similarity, it may be inferred that OSC-IVM rescued oocytes have an improved degree of cytoplasmic maturation compared to oocytes matured in the Media-IVM condition, which is an important aspect for measuring the clinical utility of these rescued oocytes. Oocytes matured in co-culture with OSCs also showed minimal evidence of gene expression signatures involved in DNA damage response, DNA repair, or oxidative stress, indicative of their good health. While understanding the precise mechanism of action of OSC improvement of rescue IVM outcomes was not explored in this study, this work shows that improvement in rescue IVM demonstrated by primary granulosa cell co-culture and other supplements can likewise be accomplished through co-culture with stem cell-derived OSCs.^22,25,39^

The method of co-culture rescue-IVM described here integrates readily with existing procedures in ART laboratories, requiring no special equipment that is not commonly found in ART laboratories and minimal training of clinical embryology staff. Further, this OSC-IVM performs effectively in hCG triggered cycles with denuded oocytes. Additionally, the primarily paracrine interaction between the oocytes and OSCs in this system, in which direct connections and gap junctions are not formed and soluble factors are exchanged, allows for a broad range of potential culture configurations. The applicability to denuded oocytes provides an expansion of functionality over other IVM systems that require that COCs be left either partially or entirely intact. This is particularly important as modern IVF practice more often adopts intracytoplasmic sperm injection (ICSI) and oocyte cryopreservation, which requires denudation of oocytes at retrieval. Additionally, as this OSC-IVM method utilizes a single source of differentiated OSCs, this method requires no additional primary cell culture donation or customization per patient, making it a readily usable future method in IVF treatment.

## Limitations and Reasons for Caution

While the number of subjects and oocytes utilized in this study was limited, our findings represent an important first step in establishing stem cell-derived OSCs as a co-culture platform for rescue IVM. More research is needed to determine the precise mechanism of action underlying OSC-IVM to characterize the likely diverse function of the cells within the culture system. More research is needed to determine the embryo formation capacity of these OSC-IVM rescued oocytes, as well as the epigenetic health of embryos generated in OSC-IVM. In addition, future studies will determine if OSC-IVM embryos are capable of healthy implantation, development and live birth compared to traditional rescue IVM and IVF controls. Regardless of the limitations, the use of OSCs derived from a single source of hiPSCs represents a novel approach for reproducible, highly standardized delivery of ovarian support cell co-culture in IVF and rescue-IVM settings, a key practical consideration for use in a clinical treatment setting.

## Acknowledgments

This work was performed with the support of clinical partnerships at Extend Fertility of New York, RMA of New York and Ruber Clinic of Madrid. We thank the dedicated support and work of the embryology and support staff at these clinics for coordinating and managing the collaborative study. We thank the Wyss Institute for Biologically Inspired Engineering at Harvard University for material transfer of reagents used in this preclinical work. We thank Professor Mary Herbert, Professor Phillip Jordan, Professor George Church, Professor Kristin Baldwin, and Dr. Sara Vaughn for advice and guidance on the use of OSCs in rescue-IVM work. We also thank the New York University flow cytometry and imaging cores for their assistance in analysis of rescue-IVM outcomes and OSC production. We likewise thank the teams at New England Biolabs, Abberior Inc., Illumina and Azenta Life Sciences for technical advice in oocyte analyses.

## Author Contributions

C.C.K. designed, supervised, and coordinated the study as well as provided technical assistance in embryology, cell engineering, production, qualification, and sequencing. S.P., A.G., M.M., F.B., S.L.E. and C.A. performed embryology work for the study. K.S.P., B.P., and A.D.N. produced and qualified OSC batches, performed embryology work, and performed RNA-sequencing library preparation. G.R. performed all transcriptomic and statistical analysis. A.F. coordinated study logistics. D.H.M. and K.W. assisted in study design, stimulation concepts, and embryology workflows. M.P.S., P.R.J.F., and P.C. assisted in OSC data interpretation. D.A.K., M.F., and S.M., assisted with oocyte collections and informed consent J.U.K., A.C.,, and D.O. performed all oocyte donor stimulations and retrievals. C.C.K., K.S.P., B.P., G.R., and D.H.M. wrote the manuscript with significant input from all authors.

**Figure S1:**
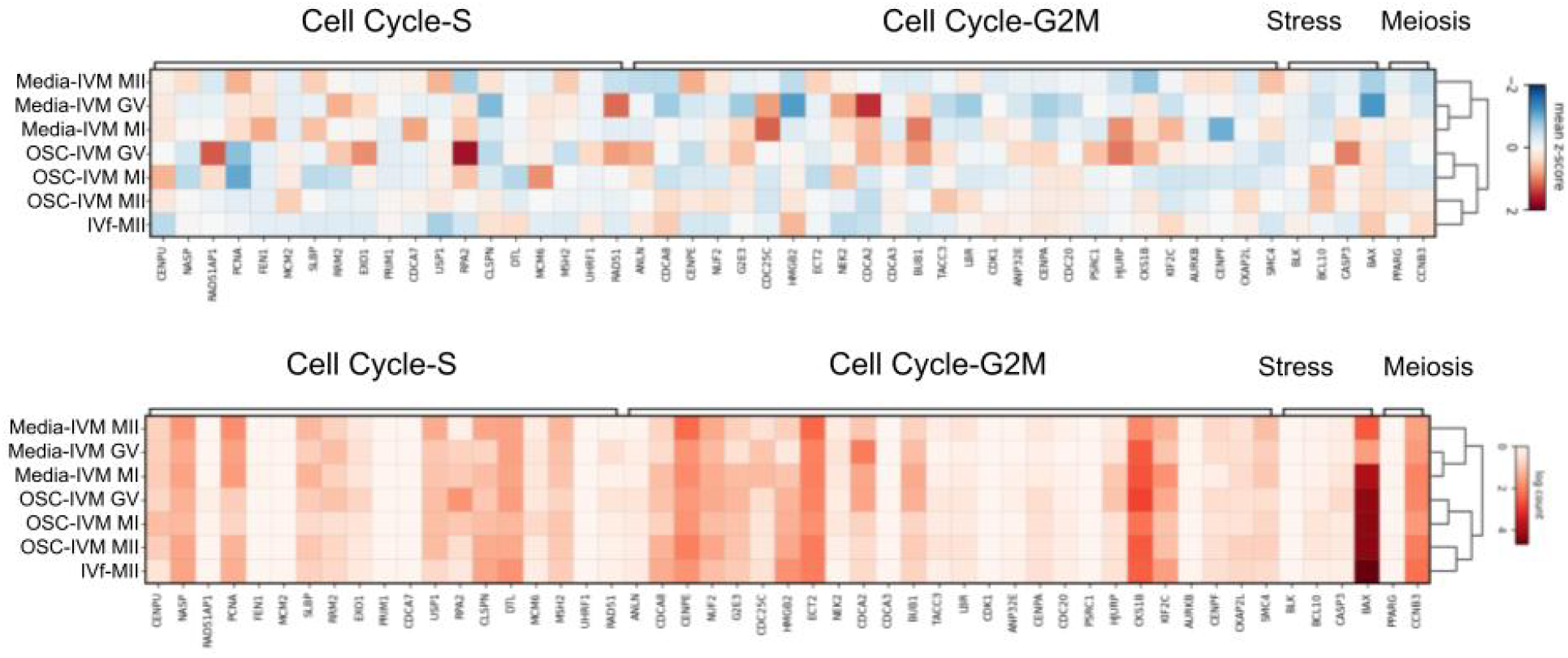
OSC-IVM oocytes show similar stress and cell cycle-related differential gene expression relative to IVF-MIIs. Gene expression values of oocytes for different developmental states (GV, MI or MII) for each experimental condition (OSC-IVM, Media-IVM, IVF-MII) are grouped for analysis, with each row representing a specific group. Relative (top panel) and absolute (bottom panel) gene expression are shown for each group for specific genes with known roles in cell cycle, stress, and meiosis, with each column indicating a specific gene. Samples are ordered on the y-axis utilizing unsupervised hierarchical clustering (UHC) for the selected genes, as a measure of relative similarity.

